# RelB proximity proteomics and CRISPR screening define chromatin regulators of noncanonical NF-κB control of HIV latency and reactivation

**DOI:** 10.64898/2026.08.11.744217

**Authors:** Cameron R. Bussey-Sutton, Carley N. Gray, Benjamin J. Wu, Jackson J. Peterson, Samuel D. Burgos, C. Allie Mills, Laura E. Herring, Edward P. Browne, Anne-Marie W. Turner, David M. Margolis, Michael Emerman, Brian D. Strahl

## Abstract

Activation of the non-canonical NF-κB pathway via RelB/p52 signaling by SMAC mimetics such as AZD5582 is a promising strategy to induce HIV expression from latency, but the chromatin mechanisms linking RelB/p52 to proviral regulation remain poorly defined. Here, we combine RelB BioID proteomics, targeted HIV-CRISPR screening, and pharmacologic validation to identify ncNF-κB-associated regulators of HIV expression. RelB BioID revealed an extensive interaction network of chromatin and transcriptional regulators in basal and AZD5582-activated states. Basally associated RelB proteins included NSD2, SWI/SNF components, UHRF1, and DNMT1, while AZD5582-enriched proteins included p300, USP7, LSD1/KDM1A, NuRD components, SIN3A, and HBO1/KAT7. Functional screening using a custom guide RNA library targeting all BioID-identified factors identified regulators that promote HIV reactivation, including HBO1/KAT7, NSD2, and SIN3A, and regulators that restrict HIV expression, including p300, CHD4, and USP7. Because KAT7/HBO1 and p300 encode acetyltransferases with opposing screen phenotypes, we tested whether their catalytic activities contribute to HIV transcriptional regulation and found that, consistent with the CRISPR screen, KAT7/HBO1 inhibition reduced AZD5582-induced reactivation, whereas p300 inhibition enhanced it. Together, these data define a resource linking the RelB-associated chromatin landscape to HIV latency and reactivation.

## Introduction

Although antiretroviral therapy (ART) durably controls HIV replication, it does not eradicate the pool of latently infected cells that can reignite infection when treatment is stopped (1,2). HIV cure efforts therefore depend on understanding how proviruses are held in a silent state and how this state can be manipulated, either by inducing viral expression for clearance or by reinforcing durable transcriptional silencing. One approach, termed “shock-and-kill,” uses latency-reversing agents (LRAs) to induce HIV transcription from latently infected cells (3,4). Because HIV latency is controlled in part by chromatin state, transcription factor availability, and RNA polymerase II elongation, many LRAs target epigenetic and transcriptional regulators, including histone deacetylases, Polycomb-associated pathways, and BET bromodomain proteins (5–10). These studies have also highlighted that combinatorial strategies are likely required to achieve robust and consistent HIV reactivation.

Recent advances in identifying effective new LRAs have uncovered inhibitor of apoptosis protein (IAP) antagonists, also known as SMAC mimetics, as potent activators of the non-canonical NF-κB pathway (11–14). AZD5582 is one such compound and has been shown to promote HIV and SIV expression through activation of RelB/p52-containing ncNF-κB complexes that target the HIV promoter (**Figure 1A**). AZD5582 is especially effective when used with other LRAs, such as BRD4 inhibitors (15,16). However, despite the promise of ncNF-κB activation as a latency-reversal strategy, the downstream chromatin and transcriptional mechanisms that connect RelB/p52 signaling to HIV transcription remain poorly defined.

**Figure 1.**
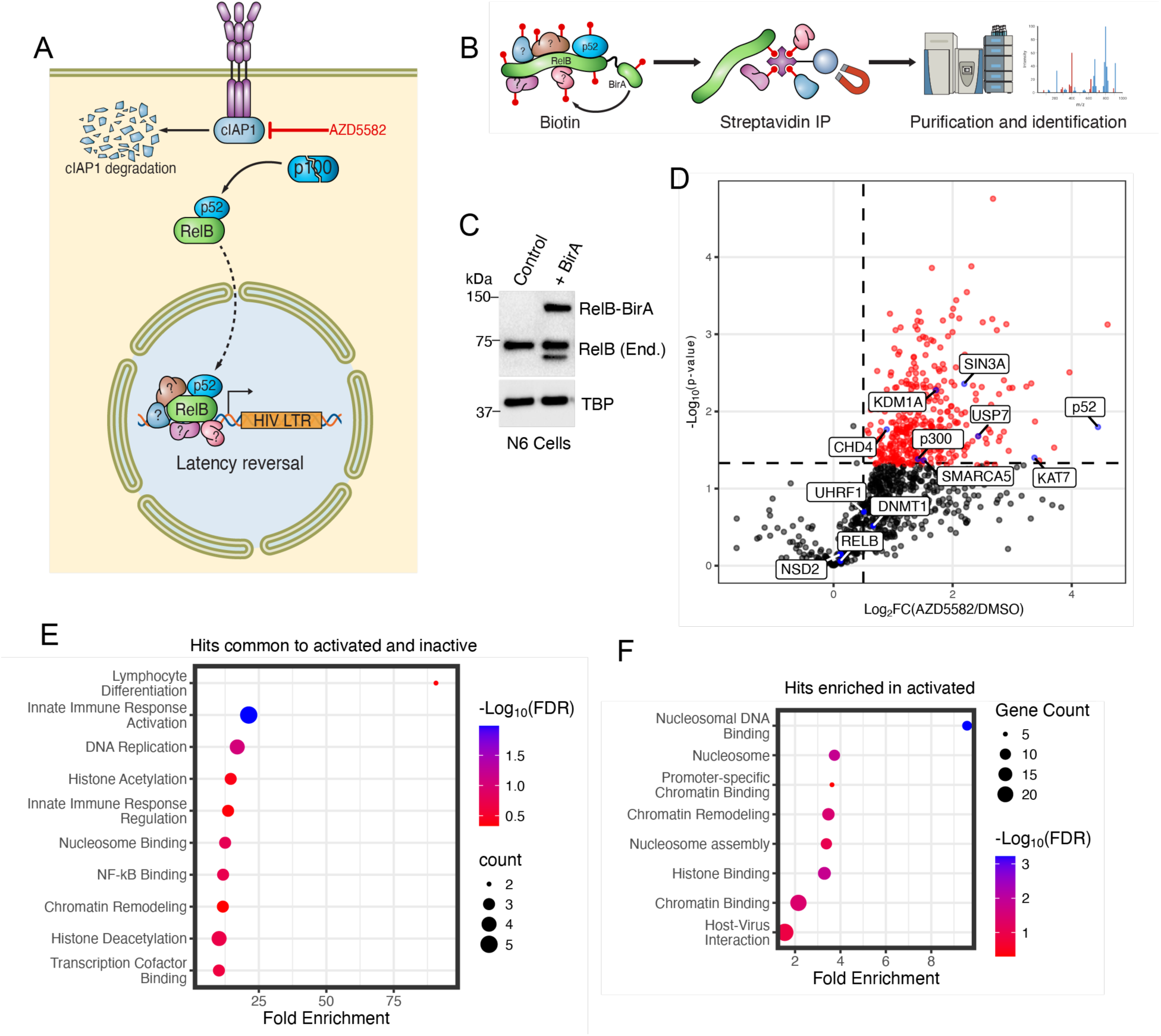
Activation of non-canonical NF-κB enriches for chromatin-modifying enzymes and regulatory factors. **(A)** Schematic highlighting the non-canonical NF-κB pathway and its role in HIV latency reversal. The antagonist of inhibitor of apoptosis proteins (IAPs), AZD5582 – a known HIV latent reversing agent (LRA) – is shown in red. Factors containing question marks surrounding RelB/p52 are used to emphasize that our understanding of how AZD5582 mechanistically functions to activate HIV transcription is not known. **(B)** Schematic of the doxycycline (dox)-inducible RelB-BirA fusion and the experimental approach of the proximity biotinylation method. **(C)** Western blot analysis showing the exogenous expression of the RelB-BirA in respect to endogenous RelB. The integrated RelB-BirA fusion, although dox inducible, was observed to express the fusion in the absence of dox in normal FBS-containing medium to levels comparable to endogenous RelB (also see Supplementary Figure S1A). **(D)** Volcano plot of AZD5582-activated versus untreated control proteomics comparison, with AZD5582-activated hits (p-value > 0.05 and log2 fold change < 0.5) highlighted. The x-axis shows enrichment (Log2FC) of proteins in AZD5582-activated condition compared to the control condition and the y-axis shows -log10(p-value) from students t-test. As shown in the plot, RelB is found in both activated and untreated samples (bottom left), whereas p52 is only identified in AZD5582-activated conditions (far right). Histone-modifying enzymes and chaperones not previously known to associate with RelB/p52 are highlighted. **(E)** Gene Ontology (GO) analysis of the proteins that were not significantly different between the AZD5582-activated and untreated conditions (log2 fold change -1 < x > 1) reveal enrichment of enrichment of immune response, chromatin, and nucleosome related terms. **(F)** Gene Ontology (GO) analysis of the proteins that were enriched in the AZD5582-activated condition compared to control (p-value > 0.05 and log2 fold change < 0.5; highlighted proteins in panel (D) reveal enrichment of chromatin and nucleosome related terms.

To address this gap, we set out to define the basal and AZD5582-activated protein interaction landscape of RelB and to functionally annotate RelB-associated factors that regulate HIV expression. Using RelB BioID proximity-labeling proteomics in a Jurkat HIV latency model, we identified a broad interaction network enriched for chromatin and transcriptional regulators in both basal and AZD5582-activated states. We then generated a custom HIV-CRISPR knockout library targeting BioID-identified factors and screened for genes that altered HIV output after AZD5582 treatment. In parallel, we performed a comparative screen using TNF-α to activate canonical NF-κB signaling, allowing us to distinguish shared regulators from factors more strongly linked to ncNF-κB-driven reactivation. Finally, we used pharmacologic inhibition of selected acetyltransferase hits to test whether their catalytic activity contributes to HIV transcriptional regulation. This integrated approach identified RelB-associated factors that promote HIV reactivation, including HBO1/KAT7, NSD2, and SIN3A, as well as factors that restrict HIV expression, including p300 and CHD4. Together, these findings define a RelB-associated chromatin and transcriptional network that functionally controls HIV latency and reactivation, and provide a resource for future mechanistic studies of ncNF-κB signaling in HIV transcriptional regulation.

## Results & Discussion

### BioID of RelB in basal and AZD5582-activated states reveals chromatin and transcriptional regulators associated with ncNF-κB

Although the SMAC mimetic AZD5582 has shown promise as an LRA, the mechanisms by which ncNF-κB stimulates HIV transcription and latency reversal (**Figure 1A)** remain incompletely understood. To further define how ncNF-κB signaling functions, we employed a proximity-based biotinylation approach (17) in which full-length RelB was cloned in-frame with miniTurbo biotin ligase and stably integrated into Jurkat N6 cells, which contain an integrated HIV-1 provirus (11) (**Figure 1B**). Expression of the RelB-miniTurbo fusion was confirmed by western blot analysis and was found to be similar to endogenous RelB when cells were cultured in standard FBS-containing medium (**Figure 1C** and **Supplementary Figure 1A**). We further confirmed that AZD5582 activated the ncNF-κB pathway in these cells, as shown by processing of the p100 NF-κB2 subunit to the active p52 isoform (**Supplementary Figure 1B**). Finally, we confirmed that the RelB-miniTurbo fusion biotinylated a range of proximal proteins upon addition of biotin (**Supplementary Figure 1C**).

To define the RelB-associated protein landscape in basal and activated states, we performed proximity labeling under control (DMSO) and AZD5582-treated conditions followed by mass spectrometry (see Methods for details). We first compared DMSO-treated RelB-miniTurbo cells with the no-biotin control to identify a basal RelB-proximal network. This analysis identified 94 proteins enriched over the no-biotin control, including known RelB- and NF-κB-associated factors such as NF-κB1/p105, NF-κB2/p100, NKRF, and RelA. The basal RelB-proximal network also included RNA-binding proteins, chromatin-associated and epigenetic regulators, and transcription-related machinery, including HNRNPU, HNRNPC, FACT, CHD4/NuRD, SWI/SNF components, MED21, TOP2A, BRD1, and STAT3 (**Supplementary Table 1**).

We next used the matched AZD5582-versus-DMSO comparison to determine how ncNF-κB activation altered the RelB-proximal proteome. This analysis classified 517 proteins detected across the DMSO and AZD5582 BioID samples as either AZD5582-enriched or detected at similar levels between the two conditions (**Figure 1C** and **Supplementary Table 1**). Using a cutoff of log2FC > 0.5 and P < 0.05, AZD5582 increased the RelB association of several chromatin and transcriptional regulators, including p300/EP300, USP7, KDM1A/LSD1, SIN3A, KAT7/HBO1, CHD4, and SMARCA5, along with the active NF-κB2/p52 species expected after non-canonical NF-κB activation (**Figure 1C** and **Supplementary Table 1**). In contrast, other factors, including NSD2, UHRF1, DNMT1, BRD1, selected SWI/SNF components, and RNA-binding and RNA-processing factors such as HNRNPU, HNRNPC, and SRSF7, were detected at similar levels before and after AZD5582 treatment (**Figure 1C** and **Supplementary Table 1**). Thus, AZD5582 does not simply create an entirely new RelB-associated proteome but instead enriches a set of chromatin and transcriptional regulators on top of a broader preexisting RelB-proximal landscape.

In contrast, other factors, including NSD2, UHRF1, DNMT1, BRD1, multiple SWI/SNF components, and RNA-binding and RNA-processing factors including HNRNPU, HNRNPC, and SRSF7, were detected at similar levels before and after AZD5582 treatment, indicating that RelB is associated with a preexisting regulatory network that is maintained during ncNF-κB activation (**Figure 1C** and **Supplementary Table 1**). Finally, Gene Ontology (GO) analysis of proteins shared between basal and AZD5582-treated states, as well as those enriched after AZD5582 treatment, revealed enrichment for immune-response, chromatin, and nucleosome-related terms (**Figure 1E–F**). These data show that RelB is associated with broad transcriptional and chromatin-modifying machinery in both basal and activated states, suggesting that ncNF-κB may engage preexisting regulatory complexes while also acquiring or enriching additional chromatin regulators after pathway activation.

### Functional annotation of RelB-associated factors using an HIV-CRISPR latency screen

To determine which of the RelB-associated factors identified by BioID were functionally important for HIV re-activation from latency, we adapted a previously described HIV-CRISPR latency screening strategy (18,19). In this system, a guide RNA library is cloned into an HIV-derived vector that can be packaged upon proviral reactivation, allowing guide abundance in viral supernatants to serve as a readout for gene knockouts that increase or decrease HIV production. We generated a custom guide RNA library targeting BioID-identified factors, including proteins associated with RelB under basal conditions, after AZD5582 treatment, or in both states. In total, this RelB-focused library targeted 517 genes with six guides per gene and included 163 non-targeting control guides, for a total of 3,265 guides. The pooled guides were cloned into an HIV-CRISPR vector containing two functional LTRs, a packaging signal, guide RNA cassette, and Cas9 (20). We then transduced J-Lat 10.6 cells, a Jurkat T-cell latency model containing a near-full-length latent HIV provirus (21), with the RelB-focused HIV-CRISPR library and treated the cells with AZD5582 to induce HIV reactivation.

In this HIV-CRISPR screen, knockout of a gene can enhance proviral reactivation, reduce proviral reactivation, or have little to no effect. Genes whose loss enhances reactivation are expected to show enrichment of their corresponding guide RNAs in the viral supernatant, whereas genes whose loss impairs reactivation are expected to show guide depletion. To quantify these effects, guides were sequenced from both viral supernatant and input cell populations, and enrichment scores were generated using MAGeCK, which accounts for the behavior of multiple guides targeting each gene (22). Analysis of J-Lat 10.6 cells transduced with the RelB-focused HIV-CRISPR library and treated with AZD5582 identified genes whose knockout either increased or decreased HIV output relative to non-targeting controls (**Figure 2A**). Bolded hits in **Figure 2** correspond to factors highlighted in the RelB BioID dataset in **Figure 1C**.

**Figure 2.**
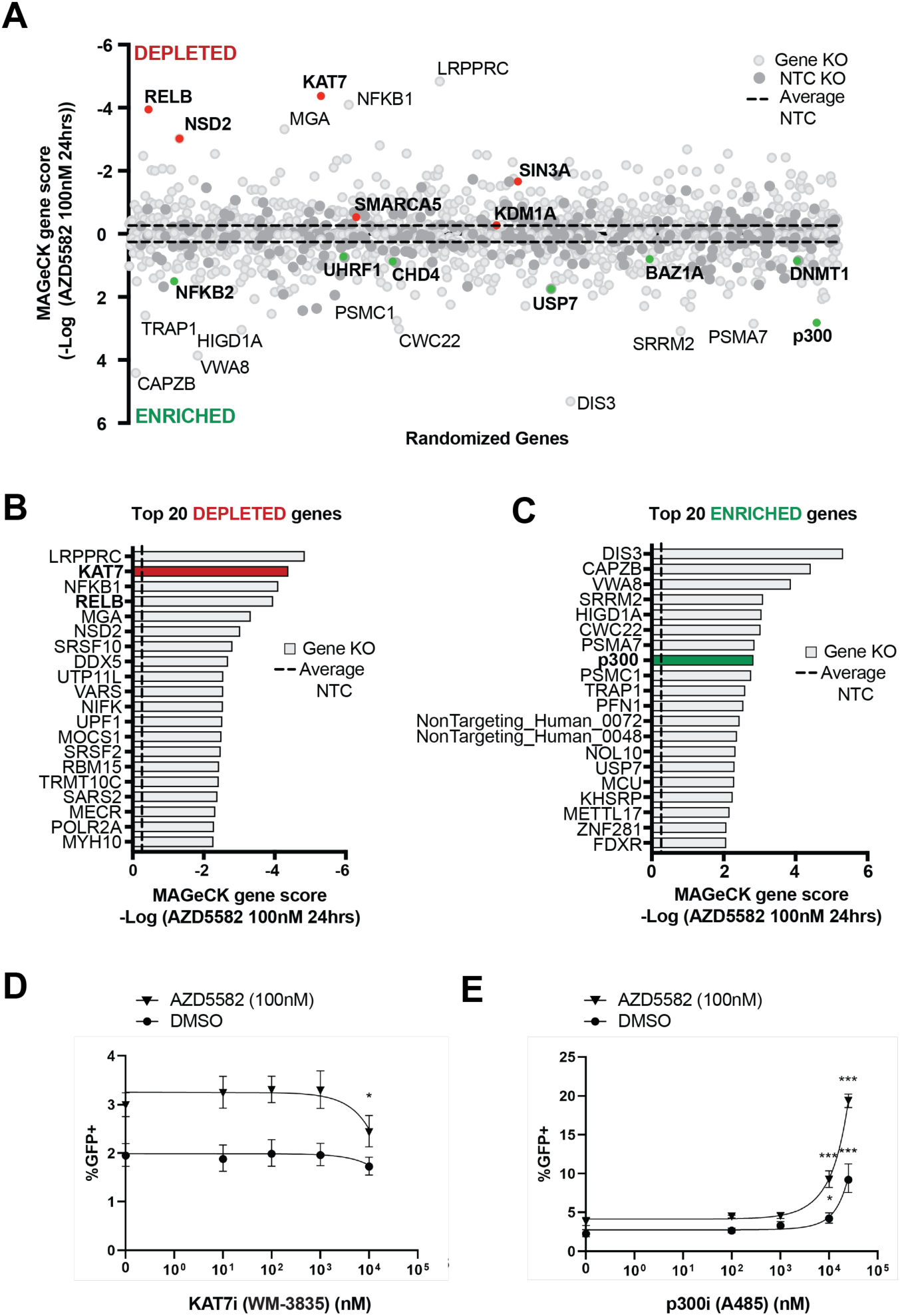
CRISPR screen identifies chromatin regulators that enhance or impair HIV reactivation downstream of AZD5582. **(A)** CRISPR knockout screen employiog a guide library of factors identified in the Bio-ID screen (see Figure 1). Upon reactivation with AZD5582, guides are packaged inside of virions and the ratio of the guides detected in the supernatant vs in the cell can be used to calculate if a guide is depleted or enriched shown as the -Log MAGeCK gene score. Gene knockouts that are enriched are predicted to improve reactivation with AZD5582, and gene knockouts that are depleted are predicted to prevent reactivation with AZD5582. Bolded genes were highlighted in Figure 1. **(B-C)** The top 20 depleted (B) or enriched (C) genes in the AZD5582 screen. Bolded genes correspond to those highlighted previously in A. Red or green bars highlight genes chosen for validation. **(D-E)** 24 hour pre-treatment with **(D)** KAT7 inhibitor WM-3835 inhibitors or **(E)** p300 inhibitor A-485, followed by 100 nM AZD5582 or an equivalent volume of DMSO in combination with previous treatment for an additional 24 hours. Inhibitor concentrations for KAT7 inhibitor WM-3835: 0, 10, 100, 1000, 10000 nM. Inhibitor concentrations for p300 inhibitor A-485: 0, 100, 1000, 10000, 25000 nM. Statistical significance for one-way ANOVA comparing 0 nM inhibitor concentration to all other concentrations in a respective dose curve (i.e. AZD5582 or DMSO) are indicated as follows: *P < 0.05; **P < 0.001; ***P < 0.0001.

Guides targeting genes in the RelB-focused library showed differential effects on HIV-transcription and as they were both depleted and enriched in the viral supernatant after AZD5582-induced reactivation of the latent provirus (**Figure 2A**). Genes on the depleted side of the screen are predicted to encode factors required for AZD5582-induced HIV reactivation, since knockout of these genes reduces viral output. As expected, RELB knockout was depleted from the supernatant, consistent with the requirement for RelB in ncNF-κB-mediated activation of HIV(23). As an additional control, loss of NFKB1 also impaired reactivation, consistent with known crosstalk between canonical and non-canonical NF-κB pathways. Among the depleted hits, we identified several chromatin and transcriptional regulators required for AZD5582-induced latency reversal, including KAT7/HBO1, NSD2, and SIN3A.

KAT7/HBO1 is a MYST-family lysine acetyltransferase that mediates H3K14ac and H4 N-terminal acetylation, which are modifications linked to transcriptional activation and other DNA-templated processes (24). Because KAT7/HBO1 emerged as a RelB-associated factor and a depleted hit in the AZD5582 screen, we tested whether its catalytic activity contributes to HIV reactivation. Pharmacologic inhibition of KAT7/HBO1 with WM-3835 partially reduced AZD5582-induced latency reversal in J-Lat 10.6 cells as well as in another latency model, N6 (**Figure 2D** and **Supplementary 2A**). Consistent with its established activity, WM-3835 reduced H3K14ac at doses that also suppress AZD5582-induced HIV transcription in the functional assays (**Supplementary Figure 2C**). These data support KAT7/HBO1 as an activating acetyltransferase pathway that contributes to ncNF-κB-driven HIV reactivation.

NSD2, an H3K36 methyltransferase (25), also emerged as a prominent depleted hit, suggesting that chromatin regulatory mechanisms beyond acetylation contribute to ncNF-κB-driven HIV reactivation. SIN3A, a scaffold of the SIN3 histone deacetylase complex (26), was also required for efficient reactivation, indicating that factors typically associated with repression may also support transcriptional responses in this context. In contrast, SMARCA5 and KDM1A/LSD1 were also found on the depleted side of the screen but had more modest effects closer to the non-targeting control distribution. These findings suggest that although many chromatin regulators associate with RelB after AZD5582 treatment, only a subset are required for HIV reactivation in this assay. A full list of screen results is provided in **Supplementary Table 2**, and the top 20 depleted genes are shown in **Figure 2B**. The strongest depleted hit was LRPPRC, an RNA-binding protein implicated in RNA metabolism and trafficking (27,28), raising the possibility that post-transcriptional RNA regulatory pathways also influence ncNF-κB-driven HIV output. Additional studies will be needed to define the role of LRPPRC and other non-chromatin hits in HIV regulation.

In contrast to depleted hits, genes enriched in the viral supernatant are predicted to encode negative regulators of HIV reactivation, since knockout of these genes increase viral output after AZD5582 treatment (**Figure 2A**, positive values). Nearly half of the BioID-identified factors included in the CRISPR library were found on the enriched side of the screen, including p300, NFKB2, CHD4, and BAZ1A. The top 20 enriched genes are shown in **Figure 2C**. Loss of CHD4, a component of the NuRD histone deacetylase complex (29), is consistent with a role for chromatin repression in limiting HIV transcription. Additional enriched candidates present in both the RelB BioID and CRISPR datasets included USP7, UHRF1, and DNMT1, factors collectively linked to ubiquitin signaling, DNA methylation, epigenetic maintenance, and transcriptional repression (30–32). In the BioID dataset, USP7 was enriched with RelB following AZD5582 treatment, whereas UHRF1 and DNMT1 were detected in the basal RelB-associated network (**Supplementary Table 1)**. Of note, recent studies support a role for USP7 and UHRF1 in HIV transcription and latency regulation (33–35). Further work will be needed to define mechanistically how these epigenetic regulators influence HIV transcription.

The recovery of p300 as an enriched hit was notable, as recent work has shown that CBP/p300 lysine acetyltransferases can inhibit HIV expression in latently infected T cells (36). Our findings agree with and extend these observations by identifying p300 as an AZD5582-enriched RelB-associated factor whose loss enhances AZD5582-induced HIV reactivation. Because p300 is a lysine acetyltransferase, we tested whether its catalytic activity contributes to HIV transcriptional regulation in this setting. Pharmacologic inhibition of p300 with A485 enhanced AZD5582-induced HIV reactivation in J-Lat 10.6 cells as well as in the N6 latency model, consistent with the CRISPR screen (**Figure 2E** and **Supplementary Figure 2B**). Consistent with its established activity, A485 reduced global H4 acetylation and H3K27ac at doses that also enhanced AZD5582-induced HIV transcription in the functional assays (**Supplementary Figure 2D**). These data support p300 as a RelB-associated acetyltransferase pathway that restricts AZD5582-induced HIV reactivation.

To distinguish factors that preferentially regulate ncNF-κB-driven HIV reactivation from those that more broadly influence NF-κB-dependent HIV expression, we performed a parallel HIV-CRISPR screen using TNF-α to activate canonical NF-κB signaling and compared the results with the AZD5582 screen (**Figure 3A–B**, **Supplementary Figure 3,** and **Supplementary Table 3**). As expected, RELB knockout strongly impaired AZD5582-induced reactivation but did not show the same depletion in the TNF-α screen, consistent with the selective requirement for RelB in ncNF-κB-mediated activation (23). In contrast, KAT7/HBO1 fell closer to the diagonal between the two screens, suggesting that KAT7/HBO1 or KAT7-associated complexes may support HIV output following both AZD5582 and TNF-α stimulation. However, acute pharmacologic inhibition of KAT7/HBO1 with WM-3835 reduced AZD5582-induced reactivation but did not suppress TNF-α-induced GFP expression under the conditions tested, instead producing a small increase at higher concentrations (**Figure 3C**). Thus, while the genetic screen identifies KAT7/HBO1 as a shared functional hit, the pharmacologic data indicate that KAT7/HBO1 catalytic inhibition has a more evident inhibitory effect in the setting of AZD5582-mediated ncNF-κB activation.

**Figure 3.**
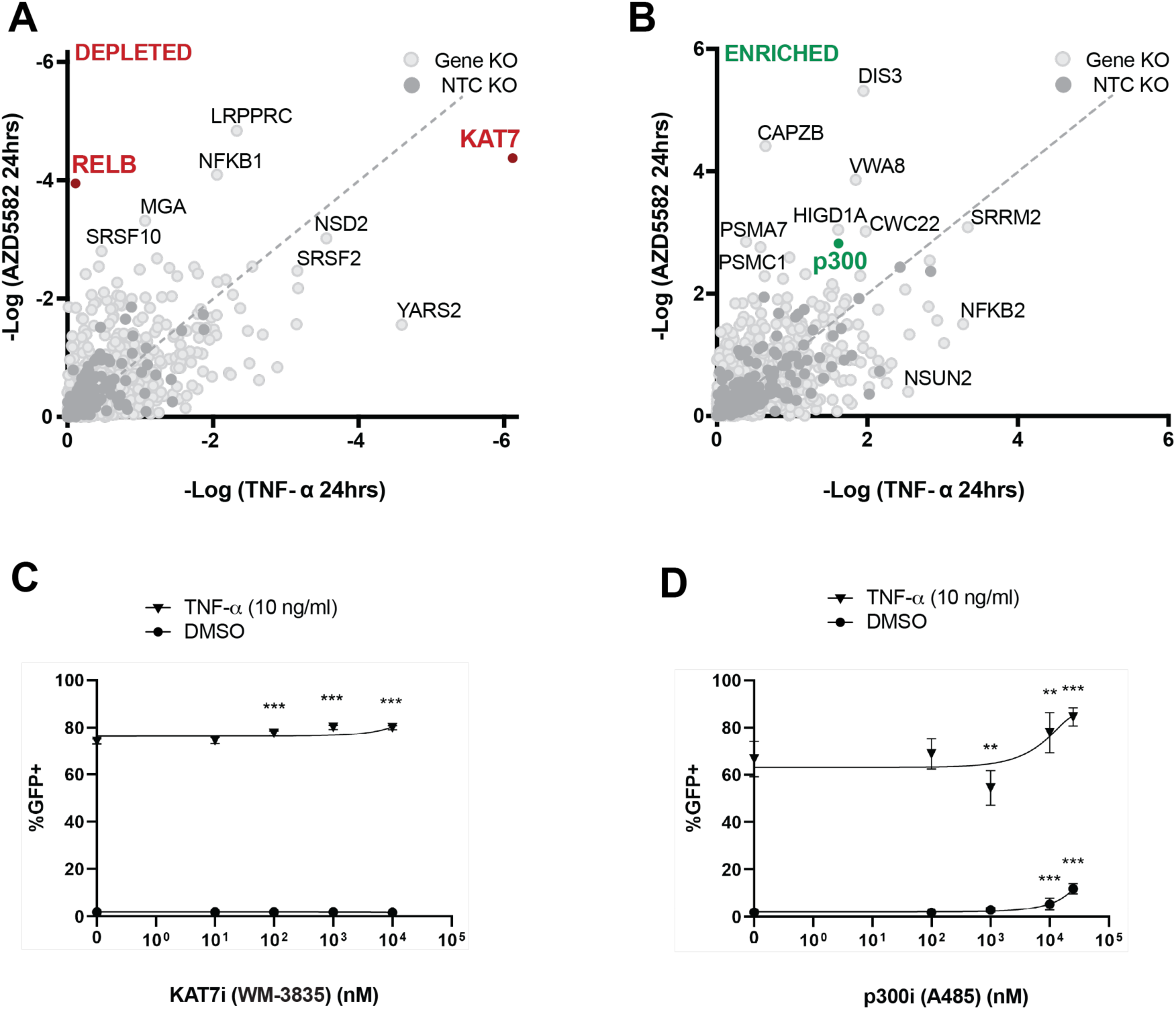
Comparative HIV-CRISPR screening distinguishes canonical and noncanonical NF-kB-associated regulators of HIV reactivation. The HIV-CRISPR screen shown in Figure 2 was compared with a parallel screen performed using the same guide RNA library and TNF-α stimulation to activate canonical NF-kB signaling. This comparison was used to distinguish genes preferentially affecting AZD5582-mediated ncNF-κB activation from those more broadly affecting NF-κB-dependent HIV expression. **(A-B)** Comparison of depleted (A) or enriched (B) genes in the AZD5582 and TNF-α screens. Select genes of interest are highlighted. In (A), genes above the diagonal show stronger depletion in the AZD5582 screen, genes along the diagonal show similar depletion in both screens, and genes below the diagonal show stronger depletion in the TNF-α screen. In (B), genes above the diagonal show stronger enrichment in the AZD5582 screen, genes along the diagonal show similar enrichment in both screens, and genes below the diagonal show stronger enrichment in the TNF-α screen. **(C–D)** J-Lat 10.6 cells were pre-treated for 24 hours with increasing concentrations of the KAT7/HBO1 inhibitor WM-3835 (C) or the p300 inhibitor A-485 (D), followed by treatment with 10 ng/mL TNF-α or an equivalent volume of DMSO in combination with the same inhibitor concentration for an additional 24 hours. HIV reactivation was measured by flow cytometry as the percentage of GFP-positive cells. WM-3835 did not decrease TNF-α-induced HIV reactivation and produced a slight increase at higher concentrations (C). A-485 increased TNF-α-induced HIV reactivation, although this effect was less pronounced than observed with AZD5582 stimulation. WM-3835 was tested at 0, 10, 100, 1,000, and 10,000 nM. A-485 was tested at 0, 100, 1,000, 10,000, and 25,000 nM. Statistical significance was determined by one-way ANOVA comparing each inhibitor concentration with the 0 nM condition within the respective dose curve. *P < 0.05; **P < 0.01; ***P < 0.001; ****P < 0.0001.

Comparison of enriched hits also revealed pathway-biased effects. For example, p300 knockout enhanced AZD5582-mediated reactivation more strongly than TNF-α-mediated reactivation, suggesting that p300 imposes a more prominent restriction on HIV expression during ncNF-κB activation. Consistent with this interpretation, pharmacologic inhibition of p300 enhanced AZD5582-induced HIV reactivation (**Figure 2E**), with a less pronounced effect observed under TNF-α stimulation (**Figure 3D**). Together, these comparative screens distinguish factors that are broadly required for HIV reactivation from those whose effects are more strongly linked to AZD5582-driven ncNF-κB signaling, providing a framework for prioritizing pathway-biased and shared regulators of HIV transcriptional control.

In summary, these studies define a RelB-associated protein interaction landscape and functionally annotate candidate regulators of ncNF-κB-driven HIV transcription. Using RelB proximity labeling, we identified a broad set of proteins associated with RelB before and after AZD5582 stimulation, including RNA-processing factors, transcriptional regulators, chromatin remodelers, histone-modifying enzymes, and epigenetic maintenance factors. By coupling this resource to a targeted HIV-CRISPR screen and pharmacologic validation of selected acetyltransferase hits, we distinguished RelB-associated factors that promote HIV reactivation, such as KAT7/HBO1, NSD2, and SIN3A, from factors that restrict HIV expression, including p300 and CHD4, while also identifying additional candidate repressive regulators such as USP7, UHRF1, and DNMT1. These findings indicate that ncNF-κB signaling is not simply coupled to transcriptional activation but instead engages both activating and repressive regulatory arms that together influence proviral output. However, it is important to note that the functional assays were focused on HIV output. Thus, the roles of RelB-associated factors identified here may differ at endogenous ncNF-κB-regulated genes and could explain why some factors showed little or no phenotype upon loss in this assay. The comparative TNF-α screen further separates regulators shared across NF-κB pathways from factors more strongly linked to AZD5582-driven ncNF-κB activation. As increasing evidence suggests that effective HIV cure strategies will require rational combinations of LRAs and/or latency-enforcing approaches, this dataset provides an important resource for prioritizing chromatin and transcriptional regulators for future mechanistic and therapeutic studies.

## Acknowledgements

This research is based in part upon work conducted using the UNC Proteomics Core Facility, which is supported in part by NCI Center Core Support Grant (2P30CA016086-45) to the UNC Lineberger Comprehensive Cancer Center, DP1 DA051110 (ME), CARE 1UM1-A1-164567 (ME, BDS, EPB, and DM), and the Genomics and Bioinformatics, Shared Resource, RRID:SCR_022606, of the Fred Hutch/University of Washington Cancer Consortium (P30 CA015704). We thank Ashokkumar Manickam for technical assistance and thoughtful comments on the study.

## Data availability

Raw and unprocessed mass spectrometric data is available at the PRoteomics IDEntifications (PRIDE) database (https://www.ebi.ac.uk/pride/).

## Author contributions

C.R.B.-S. and B.D.S. conceived the project. C.N.G. performed the HIV-CRISPR screens and designed the guide libraries. C.A.M. and L.E.H. performed the mass spectrometry analyses. C.R.B.-S. and B.W. performed the BioID experiments, inhibitor validation studies, and western blot analyses. J.J.P., A.M., S.D.B. assisted with flow cytometry experiments. A.-M.W.T. and E.P.B. provided supervision and guidance for HIV latency experiments. D.M.M., M.E., and B.D.S. supervised the project. B.D.S., C.R.B.-S., C.N.G., and M.E. wrote the manuscript with input from all authors.

## METHODS

### Cell Culture

N6 cells (11) were maintained in RPMI media with 10% fetal bovine serum (FBS) and 500 nM efavirenz and were split every 3–4 days to maintain a cell density of around 0.3–1 million cells per mL. These cells were transduced with the miniTurbo-RelB construct and maintained in RPMI media with 10% fetal bovine serum (FBS), 500 nM efavirenz, and 7.5 ug/mL blasticidin and were split every 3–4 days to maintain a cell density of around 0.3–1 million cells per mL. For J-Lat 10.6, these cells were maintained in in RPMI media with 10% fetal bovine serum (FBS) and Gibco Penicillin Streptomycin at 90 U/mL and were split every 3-4 days to maintain a cell density of around 0.1-1 million cells per mL.

### RelB Bio-ID

The miniTurbo-RelB fusion protein vector was constructed by cloning RelB into the mammalian Tet-On Inducible Gene Expression PiggyBac Vector (pXLone-HA/Tet3G-3xFLAG/miniTurbo). Human RelB cDNA (NM_006509.4) was inserted 3’ to the miniTurbo protein (37) separated by a 5 amino-acid (GGSGG) linker. 1ug miniTurbo-RelB vector was nucleofected into 3×10^5 N6 Jurkat cells using a Lonza 4D nucleofector device (buffer SE, program CL-120). Two days after nucleofection, cells were selected for two weeks with 7.5 µg/mL blasticidin selection against the resistance gene (BSD) in the piggybac vector. Jurkat N6 miniTurbo-RelB fusion cells were treated with 100nM AZD5582 or DMSO for 24 hours. Non-nucleofected Jurkat N6 cells were also treated with the same conditions and used as controls. 30 minutes prior to harvesting, cells were pulsed with a final concentration of 50 µM biotin. Media was removed, and cells were washed twice in 10 mL of cold PBS with centrifugation at 1500 rpm for 10 minutes at 4°C. For nuclear extraction, thawed pellets were resuspended in 200 µL of Buffer A (10 mM HEPES (pH 8.0), 10 mM KCl, 0.5% NP-40, protease and phosphatase inhibitors) with 1 mM DTT and incubated at room temperature for 10 minutes. Samples were centrifuged at 15,000 × g for 3 minutes at 4°C, and the supernatant containing the cytoplasmic fraction was collected. Pellets were washed in 500 µL of Buffer A (without DTT) to remove residual cytoplasm. After a final spin at 15,000 × g for 1 minute at 4°C, supernatants were removed, and nuclei were subjected to lysis in 500 µL of RIPA+Urea lysis buffer (50 mM Tris-HCl [pH 7.4], 150 mM NaCl, 1% NP-40, 0.25% deoxycholate, 1 mM EDTA, 2M urea, supplemented with protease and phosphatase inhibitors). Cells were incubated on ice for 15 minutes to allow swelling, followed by sonication (5 pulses at 30% output). Lysates were clarified by centrifugation at 13,300 rpm for 10 minutes at 4°C, and protein concentrations were determined using BCA assay. Protein amounts were normalized across samples and adjusted to 1 mL with a lysis buffer. An aliquot (5%) was reserved as input. Streptavidin T1 Dynabeads (Thermo) were washed three times in RIPA+Urea lysis buffer, using 25 µL of beads per sample. Beads were resuspended to create a slurry, with a final volume such that 100 µL of slurry was added to each clarified lysate. Samples were incubated overnight at 4°C with rotation. The flow-through was saved, and beads were washed four times for 8 minutes each with RIPA+Urea lysis buffer, followed by three washes for 5 minutes each with 50 mM ammonium bicarbonate (ABC, pH 7.8). Beads were resuspended in 50 µL of 50 mM ABC. A 10% aliquot (5 µL) was reserved for Western blot analysis, and the remaining bead slurry was frozen and submitted for mass spectrometry analysis.

For quality check of samples sent to mass spectrometry, the samples were prepared as above, with volumes scaled. The protein lysates were separated by 3-8% Tris-Acetate SDS-PAGE and transferred to polyvinylidene difluoride (PVDF) membranes. Blots were blocked in 2.5% BSA in PBS-Triton (0.2%), then incubated overnight with Streptavidin-HRP in 2.5% BSA in PBS-Triton (0.2%). Membranes were then washed three times in TBST, followed by detection by enhanced chemiluminescence (ThermoFisher).

### Proteomics Sample Preparation

Bio-ID protein samples were subjected to on-bead trypsin digestion as previously described (38). Briefly, after the last wash buffer step during affinity purification, beads were resuspended in 50 µl of 50 mM ammonium bicarbonate (pH 8.0). On-bead digestion was performed by adding 1 µg trypsin and incubating, while shaking, overnight at 37°C. The next day, 1µg trypsin was added and incubated at 37°C for an additional 3h. Beads were pelleted, and supernatants transferred to fresh tubes. The beads were washed twice with 100 µl LC-MS grade water, and washes were added to the original supernatants. Samples were acidified by adding trifluoracetic acid for a final concentration of 2%. Peptides were dried and desalted using peptide desalting spin columns (Pierce), lyophilized and stored at -80°C until further analysis.

#### LC-MS/MS

The peptide samples were analyzed by LC/MS/MS using an Easy nLC 1200 coupled to a QExactive HF mass spectrometer (Thermo Scientific). Samples were injected onto an Easy Spray PepMap C18 column (75 μm id × 25 cm, 2 μm particle size; Thermo Scientific) and separated over a 90-minute method. The gradient for separation consisted of 5–45% mobile phase B at a 250 nl/min flow rate, where mobile phase A was 0.1% formic acid in water and mobile phase B consisted of 0.1% formic acid in ACN. The QExactive HF was operated in data-dependent mode where the 15 most intense precursors were selected for subsequent fragmentation. Resolution for the precursor scan (m/z 350–1700) was set to 60,000, while MS/MS scans resolution was set to 15,000. The normalized collision energy was set to 27% for HCD. Peptide match was set to preferred, and precursors with unknown charge or a charge state of 1 and ≥ 7 were excluded.

#### Data Analysis

Raw data files were processed using MaxQuant version 1.6.15.0 and searched against the Uniprot reviewed Human database (containing 20,396 entries, downloaded March 2021) appended with a common contaminants database (245 sequences), using Andromeda within MaxQuant. Enzyme specificity was set to trypsin, up to two missed cleavage sites were allowed, methionine oxidation and N-terminus acetylation were set as variable modifications. A 1% FDR was used to filter all data. Match between runs was enabled (5 min match time window, 20 min alignment window), and a minimum of two unique peptides was required for label-free quantitation using the LFQ intensities. Perseus was used for further processing (39). Reverse hits, and proteins with only 1 unique+razor peptide, were removed from the dataset. Proteins with >50% missing values were removed, and missing values were imputed from normal distribution within Perseus. Log2 fold change (FC) ratios were calculated using the averaged Log2 LFQ intensities of and students t-test performed for each pairwise comparison, with p-values calculated. Proteins with significant p-values (<0.05, Student’s T-test) and Log2 FC >1 were considered biological interactors. Gene Ontology and enrichment analyses were conducted using DAVID (40). Genes were searched against the default databases, as well as HIV_INTERACTION, HIV_INTERACTION_CATEGORY and HIV_INTERACTION_PUBMED_ID. GO terms used are described in Supplementary Table 1.

### Western blot analyses and antibodies

For confirmatory western blots, samples were prepared as above and after overnight incubation, beads were washed four times for 8 minutes each with RIPA+Urea lysis buffer. Beads were then incubated in 40 µL of elution buffer (30mM Biotin dissolved in 2% SDS) for 30 minutes at room temperature with intermittent mixing. To prepare samples for Western blot, 10 µL of 5x SDS loading dye + BME was added, the mixture was briefly vortexed, and the samples were boiled for 10 minutes. Magnetic beads were removed, and the supernatant was loaded onto a gel for electrophoresis. The input and flow-through lysates were prepared similarly with the addition of SDS loading dye + BME, followed by 5 minutes of boiling before loading onto a gel. The protein lysates were separated by 3-8% Tris-Acetate SDS-PAGE and transferred to polyvinylidene difluoride (PVDF) membranes. Blots were blocked in 5% milk in Tris-Buffered Saline (TBS), then incubated overnight with primary antibodies diluted in TBS containing 5% milk and washed with 0.1% Tween20 TBS (TBST). Membranes were then stained with secondary antibodies (1:10,000) conjugated to horseradish peroxidase for 1h at room temperature and washed three times in TBST, followed by detection by enhanced chemiluminescence (ThermoFisher). Antibodies used were anti-RelB (Cell Signaling and Technology 4922; 1:1000), anti-TBP (Abcam ab63766; 1:1000), Anti-Alpha Tubulin (Sigma Aldrich T9026, clone DM1A; 1:1000), and p100/p52 (EMD Millipore 05-361; 1:1000).

### CRISPR Library Construction

The RelB library contains guides that target 503 genes total was previously published for use in a different context (41). The full list of RelB genes and guides can be found in Supplement 5 of Gray et al. (41). In brief, we generated 6 guides/gene using CHOPCHOP (42), Guides (43), and Synthego gRNA algorithms and supplemented the library with 163 non-targeting controls (NTCs) which amounts to 3181 guides total in the RelB library that we ordered from Twist BioSciences San Francisco, CA. DNA sequences corresponding to each gRNA in the library was cloned into the HIV-CRISPR vector as previously described (20). Briefly, BsmBI overhangs were PCR amplified on the guide library and cloned into the BsmBI-digested (New England Biolabs, R0134S) HIV-CRISPR vector by Gibson Assembly (New England Biolabs, E2611S). To ensure guides were equally represented in the library population after cloning, next generation sequencing was performed.

### HIV-CRISPR screening in J-LAT 10.6 cells

The HIV-CRISPR screen was performed as previously described (41). Briefly, we generated lentivirus from our HIV-CRISPR library and transfected J-lat 10.6 cells (knocked out for Zinc Antiviral Protein (ZAP)) at a <1 MOI. We transduced with 500x coverage and then selected for transduced cells for 10-14 days with puromycin so that we have a pooled population of our RelB library knockout cells. For reactivation, we calculated for 3600x coverage and used 100nM AZD5582 (MedChemExpress) or 10 mg/mL of TNF-alpha (Peprotech, Cranbury, NJ, USA, 300-01A) for 24 hrs and then collected the cells and the supernatant. After extracting gDNA from cells and RNA from the supernatant and prepping ends for sequencing via PCR, we performed 50 bp sequencing on the MiSeq V3 platform by the Fred Hutchinson Cancer Research Center Genomics Shared Resources. Analysis was performed and guide enrichment or depletion was determined using the MAGeCK statistical package.

### Flow Analysis

J-Lat 10.6 cells were seeded at a density of 150,000 cells per well in a total volume of 200 μL (0.75 million/mL) in 96-well plates. Cells were pre-treated with the indicated inhibitors for 24 hours, then 100 nM AZD5582, 10 ng/mL TNF-alpha, or an equivalent volume of DMSO was added for an additional 24 hours of treatment before sample preparation for flow cytometry and flow cytometry evaluation for GFP expression. Technical replicates for sample preparation were done in quadruplicates. Each experiment was performed three times separately for three biological replicates.

N6 cells were seeded at a density of 60,000 cells per well in a total volume of 200 μL in 96-well plates. Cells were treated with either 100 nM AZD5582 or an equivalent volume of DMSO. To assess potential synergy with AZD5582, A485 (p300 inhibitor) and WM-3835 (KAT7 inhibitor) were added to separate wells at the indicated concentrations. Treatments were performed in triplicates for each condition. Cells were incubated with treatments for 48 hours before analysis by flow cytometry to evaluate activation of the HSA reporter within the provirus.

#### Evaluation of GFP expression in the Jurkat Latency Model

The J-Lat 10.6 cell line, which incorporates a GFP reporter gene within the provirus, was used to assess the frequency of latently infected cells undergoing reactivation. Cells were stained with BioLegend Zombie Violet™ Fixable Viability Kit in cold PBS (Catalog Number 423113). Staining was performed in the dark at room temperature, followed by sequential washing and fixation in 4% PFA. After final resuspension in FACS buffer, samples were analyzed using a BD FACSCelesta^TM^ Flow cytometer. Data analysis of GFP expression was conducted on live, single-cell gates using FlowJo v10.10.0.

The N6 cell line, which incorporates a heat-stable antigen (HSA) cell-surface reporter within the provirus, was used to assess the frequency of latently infected cells undergoing reactivation. Cells were stained with LiveDead Fixable Violet dye (Invitrogen, catalog L34955) and rat anti-mouse CD24 (HSA)-PE antibody (BD, catalog 553262, clone M1/69) in cold PBS. Staining was performed in the dark at room temperature, followed by sequential washing and fixation in 4% PFA. After final resuspension in FACS buffer, samples were analyzed using a BD LSR Fortessa flow cytometer. Data analysis of HSA expression was conducted on live, single-cell gates using FlowJo v10.10.0.

