## Supplementary Figures for "RelB proximity proteomics and CRISPR screening define chromatin regulators of noncanonical NF-κB control of HIV latency and reactivation"

Supplementary Materials for

### Proteomic and targeted CRISPR screen reveals the chromatin and transcriptional network of noncanonical NF-κB signaling in HIV latency reversal

Cameron R. Bussey-Sutton[1,9], Carley N. Gray[2,9], Benjamin J. Wu[1,9], Jackson J. Peterson[3,4], Samuel D. Burgos[3, 4], C. Allie Mills[5], Laura E. Herring[5], Edward P. Browne[3, 4, 6], Anne-Marie W. Turner[3, 4, 6], David M. Margolis[3, 4, 6], Michael Emerman[7] & Brian D. Strahl[1, 8]*

[1]Department of Biochemistry and Biophysics, University of North Carolina, Chapel Hill, NC, USA.
[2]Department of Microbiology, University of Washington, Seattle, WA, USA.
[3]Department of Microbiology and Immunology, UNC Chapel Hill, Chapel Hill, NC, USA.
[4]UNC HIV Cure Center, UNC Chapel Hill, Chapel Hill, NC, USA.
[5]UNC Proteomics Core Facility, Department of Pharmacology, University of North Carolina, Chapel Hill, North Carolina, USA.
[6]Department of Medicine, UNC Chapel Hill, Chapel Hill, North Carolina, USA.
[7]Division of Basic Sciences, Fred Hutchinson Cancer Center, Seattle, WA, USA.
[8]Lineberger Comprehensive Cancer Center, University of North Carolina, Chapel Hill, North Carolina, USA.
[9] These authors contributed equally

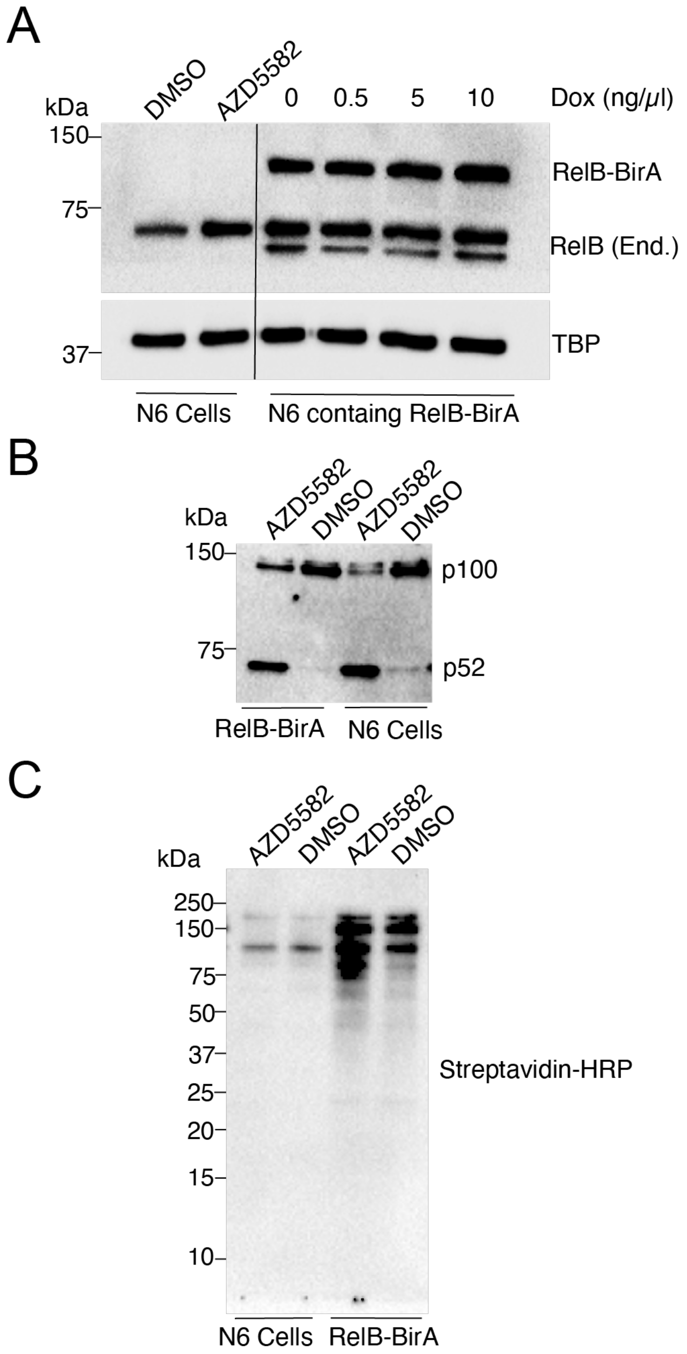


**Supplemental Figure S1 | Validation of the RelB Bio-ID system. (A)** Western blot demonstrating the expression of the RelB-BirA fusion with increasing concentrations (or without) doxycycline (dox). **(B)** Treatment of N6 cells (with or without the integrated RelB-BirA fusion) responds correctly upon addition of AZD5582 to stimulate the ncNF-κB signaling pathway, as demonstrated by the cleaving of p100 to activated p52. (**C)** Stimulation of ncNF-κB by AZD5582 results in a range of biotinylated proteins in the Bio-ID assay, as shown by streptavidin-HRP western blot analysis.

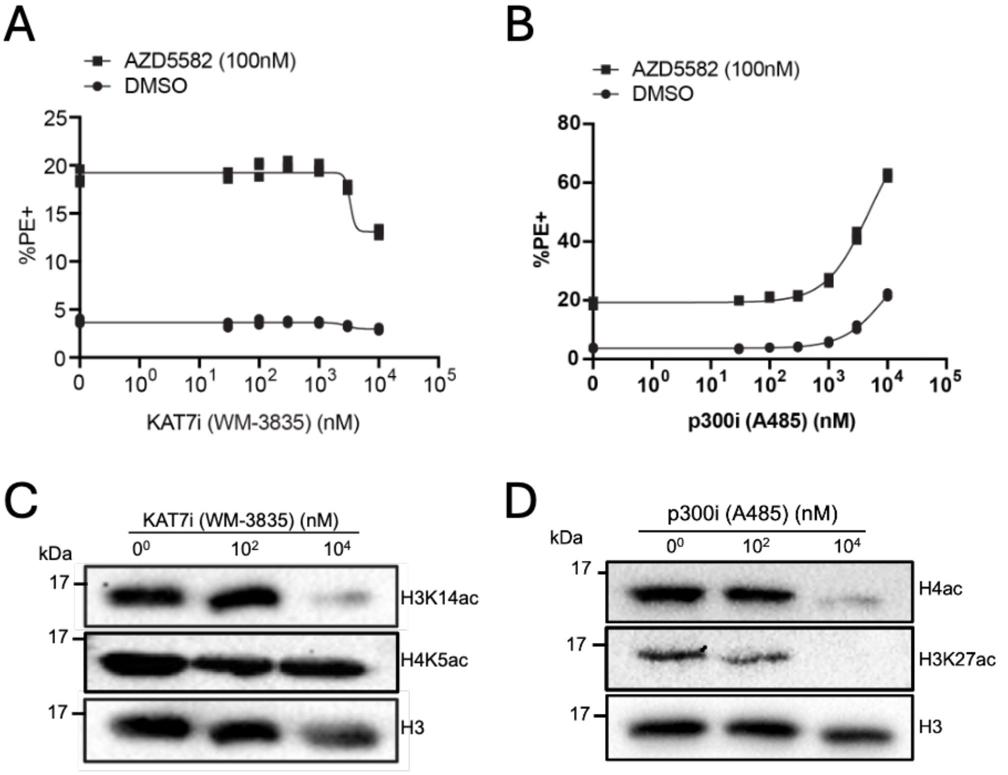


**Supplementary Figure S2 | Pharmacologic and western blot validation of KAT7/HBO1 and p300 inhibition in the N6 latency model. (A-B)** N6 cells were treated for 48 hours with increasing concentrations of the KAT7/HBO1 inhibitor WM-3835 (A) or the p300 inhibitor A485 (B) and 100 nM AZD5582 or an equivalent volume of DMSO. HIV reactivation was measured by flow cytometry as the percentage of PE-positive cells. WM-3835 partially reduced AZD5582-induced HIV reactivation, whereas A485 enhanced AZD5582-induced HIV reactivation. (C-D) Western blot analysis of N6 cells treated with increasing concentrations of WM-3835 (C) or A485 (D). WM-3835 reduced H3K14ac, consistent with inhibition of KAT7/HBO1-dependent histone acetylation. H4K5ac and total H3 are shown for comparison and loading. A485 reduced global H4ac and H3K27ac, consistent with inhibition of p300/CBP-dependent histone acetylation. Total H3 is shown as a loading control. Together, these results show that inhibitor doses affecting HIV reactivation also altered the expected histone acetylation marks.

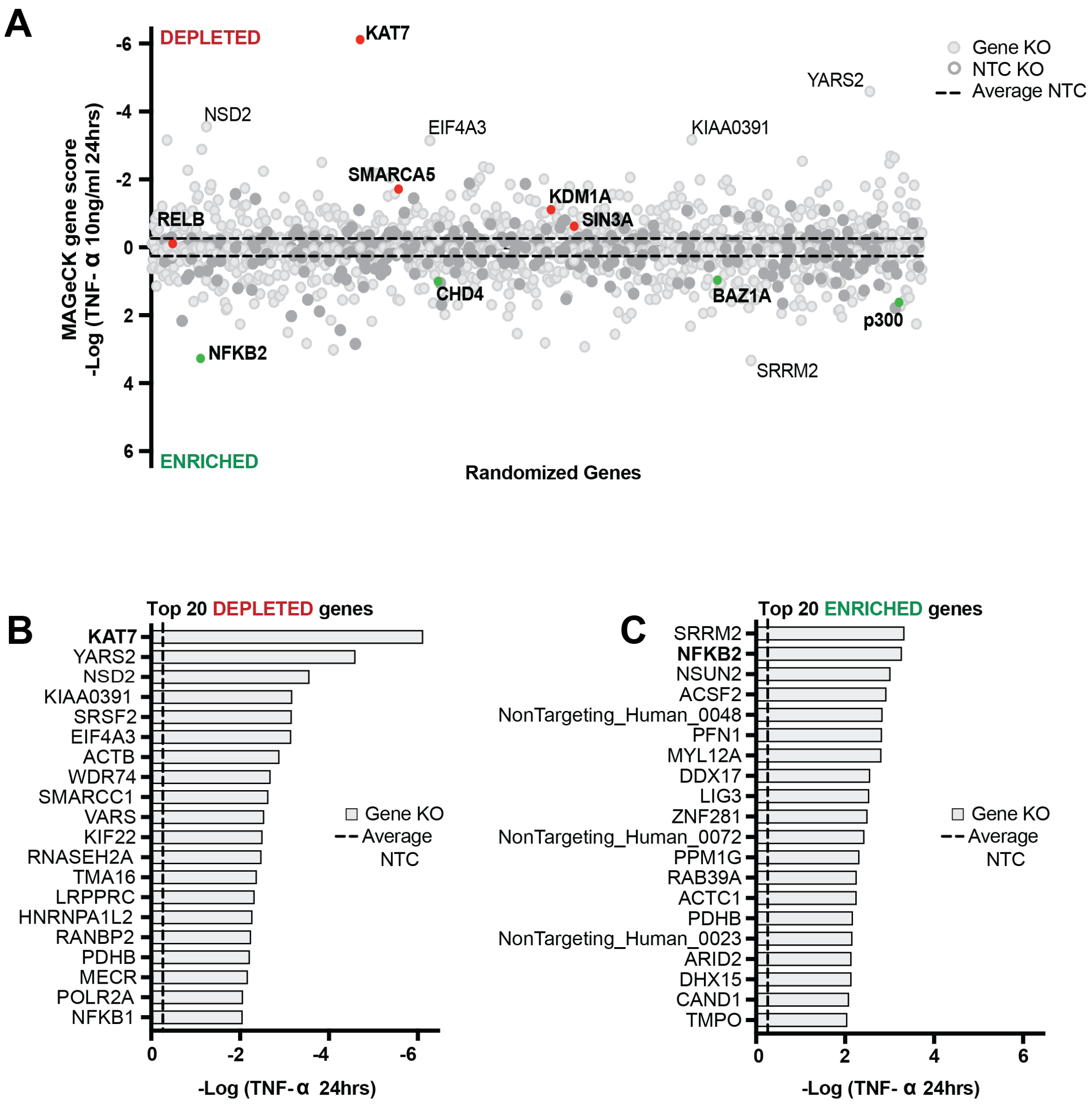


**Supplemental Figure S3 | A CRISPR screen of NF-kB targeting factors to predict the phenotype of gene knockouts and if they will improve or prevent reactivation with TNF-α** **(A)** A CRISPR screen where a pool of J-lat 10.6 cells that are knocked out for a library of NF-kB targeting factors were reactivated with TNF-α. Guides are packaged inside of virions upon reactivation and the ratio of the guides detected in the supernatant vs in the cell can be used to calculate if a guide is depleted or enriched shown as the -Log$_2$ MAGeCK gene score. Genes falling on the depleted side, knockout is predicted to prevent reactivation with TNF-α. Genes falling on the enriched side, knockout is predicted to increase reactivation with TNF-α. Bolded genes correspond to those highlighted in Figure 1D **(B-C)** the top 20 depleted (B) or enriched (C) genes.
